# Cyclin C Promotes Pancreatic Development and Suppresses Cancer Initiation Through Maintenance of the Autophagy-Lysosome Pathway

**DOI:** 10.1101/2024.08.21.609015

**Authors:** Sara E. Hanley, Kathy Q. Cai, Stephen D. Willis, David C. Stieg, Andres J. Klein-Szanto, Kerry S. Campbell, Randy Strich

## Abstract

The cyclin C (Ccnc)-Cdk8/Cdk19 kinases are components of the Mediator that represses or activates gene transcription. The present study found that Ccnc is required for both steady state and induced autophagic gene transcription in mouse embryonic fibroblasts. In vivo, pancreatic ablation of *Ccnc* (*Ccnc^PanΔ^*) resulted in phenotypes (islet atrophy, acinar cell damage) also observed in autophagy deficient models. However, *Ccnc^PanΔ^*animals displayed more dramatic phenotypes including earlier death and accelerated acinar ductal metaplasia (ADM) and pancreatic intraepithelial neoplasia (PanIN) precancerous lesion formation when these animals also expressed oncogenic *Kras^G12D^*. Consistent with the in vivo results, a *Kras^G12D^*; *Ccnc^PanΔ^* pancreatic derived cell line displayed reduced autophagy lysosome pathway (ALP) activation. Although autophagy deficient acinar cells undergo Tp53-dependent cell death, histopathology revealed that *Kras^G12D^*;*Ccnc^-/-^* (PC) pancreata did not allowing damaged cells to keep dividing. Therefore, Ccnc both supports normal ALP function protecting endocrine and exocrine cell but also kills damaged cells before they become malignant. Finally, the PC cell line displayed reduced proteasome function rendering cells hypersensitive to proteasome inhibitors. This hypersensitivity was also observed in disparate *Ccnc^-/-^* tumor cell lines or in *Ccnc^+/+^* tumor cells co-treated with Cdk8/Cdk19 inhibitors. These findings suggest a new avenue to target pancreatic neoplasms by inhibiting cyclin C-Cdk8/Cdk19 proteasome activity.

## INTRODUCTION

Pancreatic ductal adenocarcinoma (PDAC) is the most common pancreatic cancer being diagnosed in about 64,000 individuals in the U.S. in 2021 and resulting in about 50,000 deaths (Siegel et al. 2023). Prognosis is extremely poor (5-year relative survival rate of <5%) since most patients are diagnosed with advanced disease. Sporadic PDAC is often associated with KRAS activation with cells undergoing acinar-to-ductal metaplasia (ADM) leading to pancreatic intraepithelial neoplasia (PanIN) arising in these ducts. Normally, ADM lesions undergo proliferation but then re-differentiate back to functional acinar cells following repair. However, the presence of activated KRAS, the de-differentiated ADM cells are locked into a proliferation program generating pancreatic PanINs (De La et al. 2008; Habbe et al. 2008). These pre-cancerous lesions are an often-used route to PDAC disease. More advanced disease is associated with inactivation of tumor suppressors such as CDKN2/INK4A and/or p53 (Hruban et al. 2001). Chronic pancreatitis is often a preamble to PDAC by causing acinar cell damage and ADM. Acinar pathology involves inflammatory injury and can be caused by multiple agents including alcohol, genetic and/or environmental factors (Kleeff et al. 2017).

The autophagy-lysosomal pathway (ALP) is a proteolysis system that identifies and destroys various cargos including toxic protein aggregates, damaged organelles, or recycles cytoplasmic content in response to nutrient deprivation (Feng et al. 2014). Two major types of autophagy have been described. First, bulk or non-selective autophagy is induced in response to starvation and usually degrades cytoplasmic constituents to provide metabolites to maintain growth (Feng et al. 2014). Selective autophagy uses receptors that target specific cellular components including damaged organelles or protein aggregates (Grumati and Dikic 2018). In the pancreas, ALP is essential for normal ß-cell formation, insulin production (Marasco and Linnemann 2018) and to prevent many disease states (Choi et al. 2013). For example, autophagy prevents pathologies such as pancreatitis and cancer initiation (Gukovskaya et al. 2017). In murine models, disruption of autophagic genes (*Atg5* or *Atg7*) cause ADM which stimulates inflammation or even cell death (Antonucci et al. 2015; Diakopoulos et al. 2015).

Under normal conditions, cyclin C (Ccnc) and its cognate kinases Cdk8 or Cdk19, are components of the CDK8 kinase module (CKM) (Bourbon 2008) that regulates transcription through association with RNA polymerase II Mediator complex (Fant and Taatjes 2019). Recent transcriptome analysis revealed that, depending on the locus, Ccnc represses or activates the transcription of ∼3000 genes in mouse embryonic fibroblasts (MEF) (Stieg et al. 2020). Many of these genes are involved in stress responses or differentiation pathways. In addition, we discovered a second role for Ccnc that is independent of transcription. In cells subjected to oxidative damage or anti-cancer drugs, a portion of Ccnc (but not Cdk8 or Cdk19) exits the nucleus and associates with the mitochondria (Wang et al. 2015). There, Ccnc directly interacts with the fission dynamin-like GTPase Drp1 to stimulate mitochondrial fragmentation (Wang et al. 2015; Ganesan et al. 2019). In addition, Ccnc is required for intrinsic regulated cell death (iRCD) by stimulating mitochondrial outer membrane permeability (MOMP) (Wang et al. 2015) through recruitment of the pro-apoptotic protein Bax (Jezek et al. 2019). Finally, mouse knockout studies revealed that Ccnc is required for normal development with embryos arresting around day 9.5 (Li et al. 2014). Conversely, *Cdk8* deletions arrest prior to implantation while *Cdk19* null embryos grow into adulthood (Dannappel et al. 2018). Therefore, developmental roles of the CKM appear more complex than simply altering transcription.

In the present study, we found that a conditional *Ccnc* knockout in the pancreas resulted in reduced autophagy, defective islet development and premature animal death. Combining the *Ccnc* knockout with the activated *Kras^G12D^* allele resulted in a rapid acceleration of ADM and PanIN formation compared to *Kras^G12D^*activation or *Ccnc^-/-^* deletion alone. In addition, Ccnc is also required for Tp53-induced cell death These results support a model that Ccnc supports pancreatic development and suppresses pancreatic cancer initiation through maintaining autophagy and triggering cell death for cells that are damaged.

## RESULTS

### Ccnc is required for steady state and induced autophagic gene transcription

We previously determined the Ccnc-dependent oxidative stress transcriptome in mouse embryonic fibroblasts (MEFs) (Stieg et al. 2019). These RNA-seq data indicated that Ccnc is required for the steady state transcription of several genes sets including those involved in autophagy. To verify these results, RT-qPCR was performed on a subset of these genes with mRNA prepared from wild type and *Ccnc^-/-^* MEF cells under nonstress conditions. As a control, we measured the mRNA levels of *Sqstm1*/*p62*, an autophagy gene (Katsuragi et al. 2015) that we previously found not controlled by Ccnc. Genes involved in autophagosome assembly (*Becn1*, *Uvrag*, *Agt9b* and *Map1LC3B*) exhibited reduced steady state expression in the absence of Ccnc (Fig. 1A). *Agt101* was the exception for genes examined. To determine if Ccnc is also required for autophagic gene induction, cells were treated with a sub-lethal concentration of the proteasome inhibitor MG-132 (Wang et al. 2013), a potent inducer of ALP (Laussmann et al. 2011; Li et al. 2018). These studies revealed that Ccnc is also required for the induction of these autophagy genes (Fig. 1A). We repeated these experiments genes involved in autophagosome regulation (*Trp53inp1*, *Gabarap*, *LC3*, *Dram1*, *Tollip*) (Fig. 1B) or autophagosome-lysosome fusion (*Plekhm1*, *Tecpr1*, *Stx17*, *Snap29*, *Vps33a*, *Vps16*) (Fig. 1C). Steady state levels of their mRNA decreased in the absence of Ccnc, although the extent of this reduction varied. In addition, only *Plekhm1* transcript levels modestly increased following MG-132 treatment in the *Ccnc^-/-^* MEFs (Fig. 1C). These results confirmed a positive role for Ccnc in supporting both steady state and induced mRNA levels of autophagy genes.

**Figure 1.**
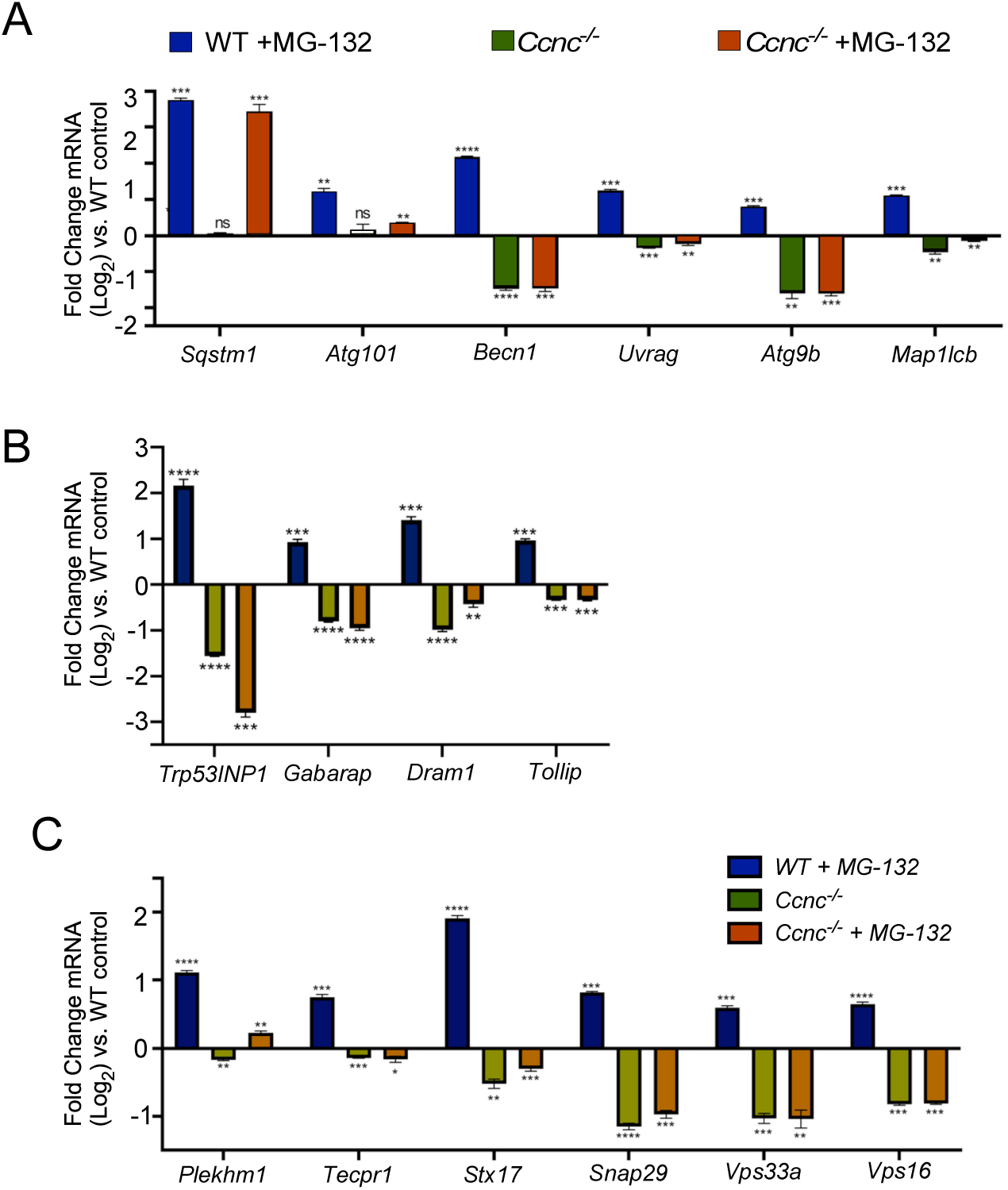
Ccnc is required for normal autophagic gene transcription. RT-qPCR analysis of mRNA levels of genes involved in autophagosome assembly (A) or autophagy regulation (B) in *Ccnc^+/+^* and *Ccnc^-/-^* MEF cells with or without MG-132 treatment (5 µM, 1 h). The horizonal line at 0 represents untreated wild-type mRNA quantitation. Values above or below this line represents the log_2_ increase or decrease in mRNA abundance, respectively. (C) RT-qPCR quantitation of mRNA of loci involved in autophagosome maturation-lysosome fusion. Asterisks indicate p values * ≤0.05, ** ≤0.01, *** ≤0.005, **** ≤0.001. Error bars = ±S.D.

### Ccnc supports normal pancreatic islet development and insulin production

Previous studies ablated the autophagy genes *Atg5* (Diakopoulos et al. 2015) or *Atg7* (Antonucci et al. 2015) in the murine pancreas and observed defective pancreatic ß-cell development, reduced insulin production and acinar cell damage. Therefore, we chose the pancreas as an organ system to provide a specific readout for ALP activity. We chose the *Pdx1*-cre mouse model system, which induces the cre recombinase in early pancreatic progenitor cells (Hingorani et al. 2003). *Pdx1*-cre mice were mated to a *Ccnc*-floxed (*Ccnc^f^*) mouse (Wang et al. 2015) generating *Pdx1*-cre;*Ccnc^f/f^* (PC) mice. Mice harboring *Atg5* or *Atg7* ablation exhibited are reduced viability although ∼40% of the animals exhibited a normal life span (Rosenfeldt et al. 2013). We found that all PC animals displayed a dramatic reduction in viability compared to control mice with a median life span of 10 weeks (Fig. 2A). By eight weeks, the PC animals all exhibited reduced weight (Fig. 2B), severe lethargy, and reached humane endpoints by 15 weeks of age. Reduced weight in the PC animals suggested the possibility of pancreatic dysfunction. Therefore, we examined islet formation by immunohistochemistry (IHC) and measured serum glucose levels in PC animals. Compared to control animals, serum glucose levels were elevated in PC animals (Fig. 2B), nearly identical to those previously described after *Atg5* ablation (Diakopoulos et al. 2015). Compared to wild-type controls (Fig. 2C), histopathology revealed dysmorphic islet appearance and reduced insulin production in PC pancreata (Fig. 2D). These results are consistent with Ccnc being required for autophagy to maintain ß-call integrity in the pancreas.

**Figure 2.**
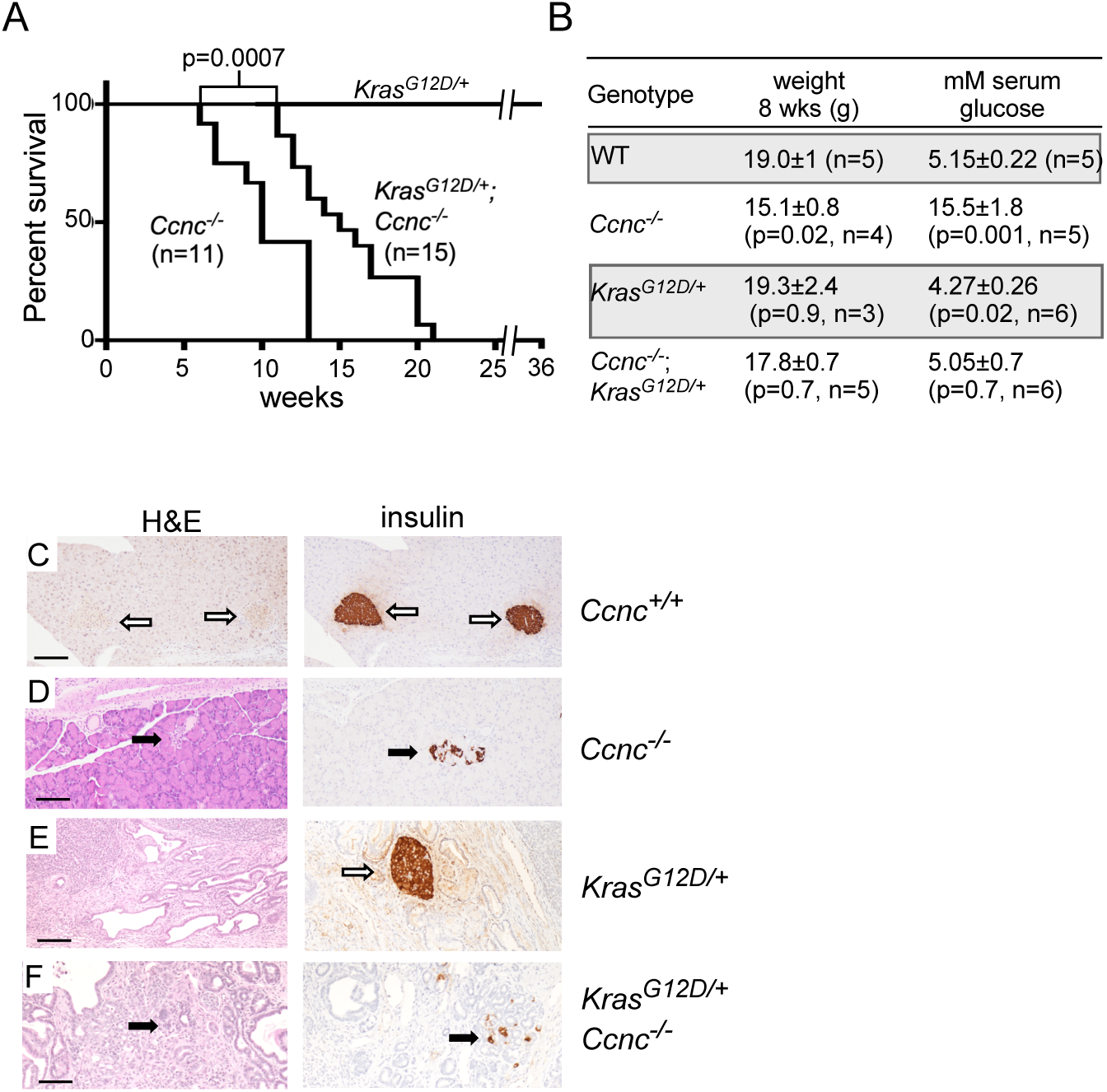
Ccnc supports pancreatic development and function. (A) Kaplan-Meier survival plot of *Pdx1*-cre;*Kras^G12D/+^* (PK), Pdx1-cre;*Ccnc^-/-^* (PC, 6:5 M:F) and *Kras^G12D/+^*;*Pdx1*-cre;*Ccnc^-/-^* (PKC, 7:8 M:F) animals as indicated. P value given for indicated survival curves. (B) Weight and plasma glucose levels are listed for eight-week-old animals with the indicated genotypes. Glucose concentrations were obtained over multiple timepoints (see Methods for details). (C-F) Corresponding images of 11- and 12-week old animals with the indicated genotypes stained with H&E or tested for insulin production via IHC. Open arrows indicate normal islet morphology and insulin production. Bars = 100 µM.

Finally, previous studies revealed that autophagy prevented acinar cell damage that caused pancreatitis or ADM (Antonucci et al. 2015; Diakopoulos et al. 2015). To test whether either result was also observed in PC animals, pancreata were obtained and analyzed by H&E staining. These studies revealed ADM in PC mice although with only partial penetrance (five out of 33 mice, ages between 5-13 weeks). The ADM lesions ranged from small focal regions (n = 2, Fig. 3A and 3B) to extensive regions covering most of the pancreas (n = 3, Fig. 3C and 3D). This result is similar to those obtained with pancreatic ablation of *Atg5* (Diakopoulos et al. 2015) indicating that Ccnc prevents ADM, but its loss alone is not sufficient to drive PanIN formation. These results indicate that Ccnc is required for normal islet formation, insulin production and animal lifespan. In addition, we observed acinar cell injury consistent with loss of ALP function.

**Figure 3.**
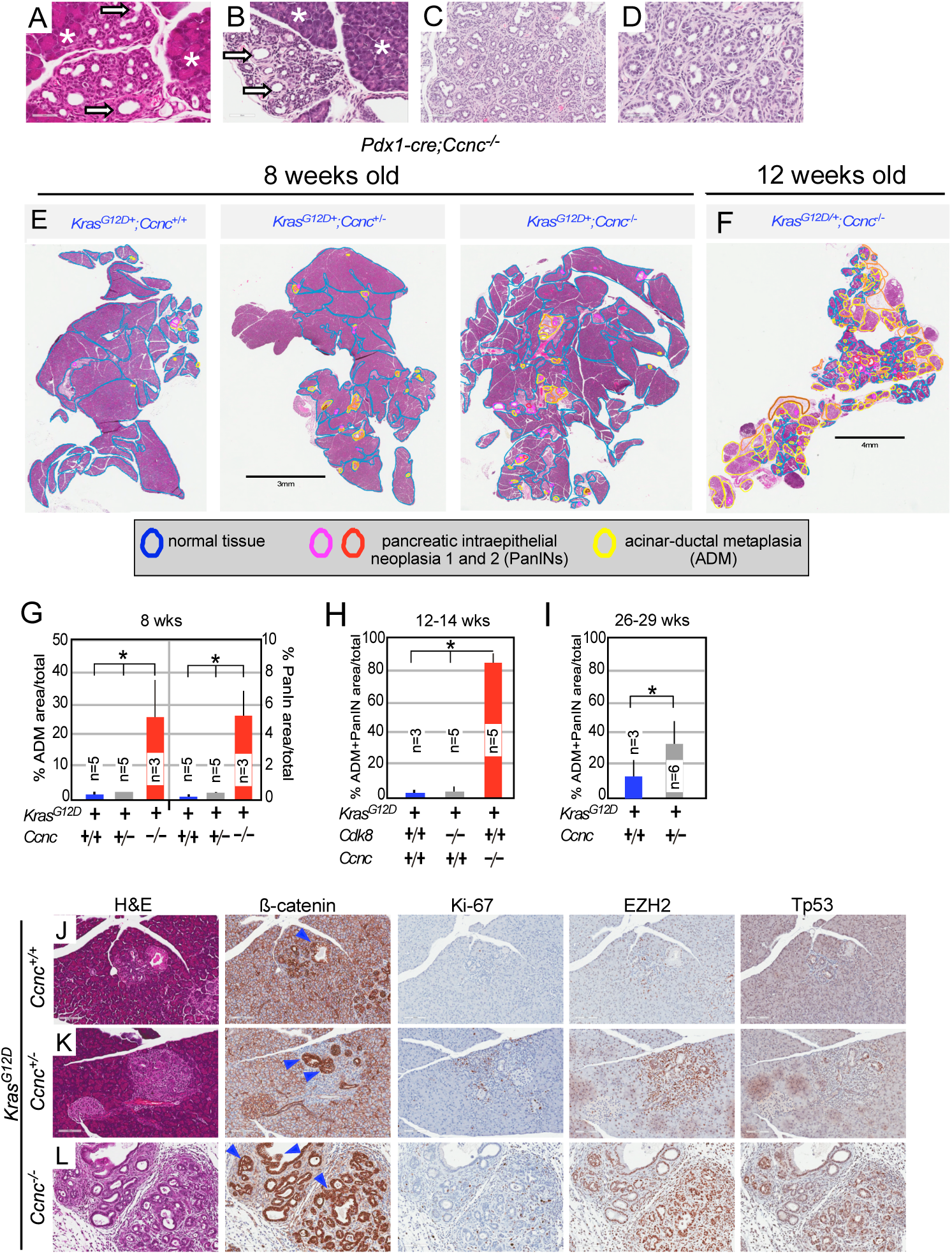
Ccnc suppresses ADM/PanIN production in the pancreas. (A-B) H&E staining of pancreatic sections from 8 wko PC. Asterisks indicate normal acinar tissues; arrows indicate ADM regions. Bars indicate 100 µM. (C-D) H&E images of extensive ADM regions in two PC animals. (E) H&E staining of pancreatic sections from 8 wko mice with the indicated genotypes. Normal, ADM and PanIN regions are indicated. (F) Histopathology of 12 wko PKC pancreas. (G) The percentage of regions with ADM and PanIN pathologies within the total pancreatic area are shown for the indicated genotypes. (H) Quantitation of ADM/PanIN areas at 12-14 or (I) 26-29 weeks for the indicated genotypes. n values for each genotype are given. Asterisks indicate statistical differences (p< 0.05). IHC analysis with the indicated antibodies of pancreatic tissues with *Kras^G12D/+^* expressed in combination with either (J) 11 week old *Ccnc^+/+^*, (K) 8 week old *Ccnc^+/-^*and (L) 8 week old *Ccnc^-/-^*) animals. Blue arrow heads indicate regions of PanIN hyperplasia. Bars indicate 100 µM.

### Oncogenic Kras^G12D^ expression stimulates PanIN production in *Ccnc* ablated animals

Constitutive activation of the *KRAS* oncogene is observed in ∼90% of PDAC and is a common feature of mouse models (Herreros-Villanueva et al. 2012). A current model posits that the presence of oncogenic Kras drives dedifferentiated ADM cells toward PanINs development (De La et al. 2008; Habbe et al. 2008). To determine if introduction of oncogenic Kras (Kras^G12D^) altered the phenotype observed in PC animals, the Lox-stop-Lox (LSL)-*Kras^G12D^* allele was introduced through mating. Cre expression removes the transcriptional “stop” DNA sequence allowing expression of *Kras^G12D^* (Hingorani et al. 2003). In contrast to PC animals, *Pdx1*-cre;LSL-*Kras^G12D^* (PK) animals did not exhibit a shortened lifespan (Fig. 2A). Consistent with this observation, PK animals displayed normal weight, serum glucose levels (Fig. 2B) and islet formation (Fig. 2E). Interestingly, the *Pdx1*-cre;LSL-*Kras^G12D^*;*Ccnc^-/-^* (PKC) double mutant animals displayed a reduced lifespan although not as severe as the PC single mutant animals (Fig. 2A). However, serum glucose levels and body weight in the double mutant PKC animals were not significantly different than the wild-type control (Fig. 2B). Surprisingly, IHC analysis revealed that the PKC animal displayed defective islet formation (Fig. 2F) as observed in the PC mutant mice. Formally, these results indicated that *Kras^G12D^* expression suppressed the elevated serum glucose phenotype observed in PC animals but only partially suppressed the shortened life span.

Next, we determined whether introduction of the activated *Kras^G12D^* allele, in combination with *Ccnc* ablation, simulated or suppressed ADM and PanIN formation. Examination of pancreata from 8-week old *Pdx1*-cre;LSL-*Kras^G12D+^* (PK) single mutant or *Pdx1*-cre;LSL-*Kras^G12D+^*;*Ccnc^f/+^* heterozygous animals revealed a modest appearance of ADM and more advanced PanIN lesions (Fig. 3E, left two panels, quantitated in Fig. 3G). The accumulation of ADM or PanIN lesions were also similar for these two genotypes at 12-14 weeks (Fig. 3H). However, a significant difference in the % of area containing ADM and PanIN-1 and PanIN-2 lesions was observed in 26-29 week old *Pdx1*-cre;LSL-*Kras^G12D+^*;*Ccnc^f/+^* animals compared to the PK control (Fig. 3I).T hese results are consistent with previous studies reporting that PK animals exhibited widespread and higher-grade ADM and PanIN lesions at 8-12 months of age (Hingorani et al. 2005).

Repeating these experiments with the PKC double mutant mice revealed an 8-fold increase in both ADM and PanIN lesions (Fig. 3E, right panel, quantified in Fig. 3G). Closer examination of these sections confirmed classical ADM and PanINs (Fig. S1A and S1B). By 12-14 weeks, nearly all of the pancreas underwent ADM and PanIN transformation (Fig. 3F), quantitated in Fig. 3H). These findings indicate that Ccnc suppresses the formation of pre-cancerous lesions in the *Kras^G12D^* pancreatic cancer model. However, adenocarcinoma formation and/or metastasis were not observed in these animals. This may be due to the relatively short life span of these animals or the proposed role for autophagy in late-stage pancreatic tumor development (see Discussion). To assess whether the suppression of pre-neoplastic lesions also required the CKM component Cdk8, we generated *Pdx1*-cre and LSL-*Kras^G12D^*;*Cdk8f/f* (PK8) animals harboring homozygous *Cdk8*-floxed alleles. Surprisingly, these animals exhibited no enhancement of in ADM or PanIN lesions compared to *Kras^G12D^* expression alone (Fig. S1C and S1D, quantified in Fig. 3H). We verified the efficient cre-directed recombination event to generate the *Cdk8* null allele in the pancreas using tissue derived genomic PCR analysis (Fig. S1E). Taken together, these results indicate *Ccnc* suppresses precancerous lesion formation in the presence of an activated *Kras* allele, but the tumor suppressor function of *Ccnc* may be independent of Cdk8 (see Discussion).

### PKC pancreata exhibit aggressive cancer markers

To confirm that the PanIN lesions described above were of ductal epithelial origin, tissue sections were probed with epithelial lineage antibody ß-catenin. PK tissues exhibited modest staining (Fig. 3J) while the *Kras^G12D^*;*Ccnc^+/-^* heterozygous animals exhibited a more aberrant phenotype including increased ADM formation (blue arrowheads, Fig. 3K). PKC pancreata exhibited strong ß-catenin staining confirming the presence of ADM and PanIN lesions (Fig. 3L). Next, we investigated the aggressiveness of the lesions by staining for the proliferation marker Ki-67. Again, the PKC pancreatic tissue exhibited higher proliferation rates than PK alone or the *Ccnc^+/-^*heterozygous sample (middle panels). Finally, acinar injury leading to pancreatitis is a major contributor of ADM and PanIN formation (Murtaugh and Keefe 2015). Previous studies reported that upregulation of the H3 lysine 27 methyltransferase EZH2 is important for the dedifferentiation program necessary for restoring normal acinar function following cerulein-induced injury (Mallen-St Clair et al. 2012). To determine if PKC pancreata exhibited this damage response, EZH2 levels were followed in PanIN regions. These studies found elevated EZH2 expression in *Kras^G12D/+^*;*Ccnc^+/-^* heterozygote and PKC homozygous samples (Fig. 3K and 3L). These results suggest that preventing cell damage may be important for Ccnc-dependent ADM/PanIN suppression. Trp53 upregulation occurs in damaged cells and can indicate the activation of the iRCD pathway (Morton et al. 2010). To examine this possibility further, we examined ADM/PanIN regions for the presence of elevated Trp53 and activated caspase 3 as a marker for iRCD. Interestingly, Trp53 staining was observed in ductal cells undergoing proliferation (as evidenced by Ki67) but no indication of caspase 3 cleavage was observed (Fig. S2). Taken together, these studies indicate that even partial loss of Ccnc activity enhances ADM and PanIN generation in the presence of an activated *Kras*. However, Trp53 upregulation does not cause growth cessation or induce cell death. These results indicate that although the damage pathways have been initiated as evidenced by ADM production, either the damage is not sufficient to induce cell death or that Ccnc plays a role in triggering this response (see Discussion).

### Autophagy is reduced in Pdx1-cre;LSL-Kras^G12D/+^;Ccnc^f/f^ cells

Autophagic flux in vivo is difficult to quantify. Therefore, to further investigate the role Ccnc plays in supporting autophagy in the pancreas, a cell line (193) was developed from pancreatic tissue derived from a 14 wko PKC animal. The cell line was validated for the presence of the *Ccnc* deletion allele (Figs. S3A, S3B) and elimination of protein expression (Fig. S3C). As a comparison, we chose another pancreatic cell line (470) derived from a 15.5 month old PK mouse that displayed extensive PanINs although PDAC lesions were not noted. Previous studies revealed a dependence of Ccnc stability on the presence of Cdk8 and Cdk19 (Barette et al. 2001; Chen et al. 2023). However, loss of Ccnc did not significantly affect Cdk8 (Fig. S3D) or Cdk19 (Fig. S3E) levels in these pancreatic cells. Finally, given the frequency at which p53 is mutated in human cancers, we determined the levels of Trp53 in these cell lines as well as a downstream effector p21. These studies revealed a modest increase in p53 and p21 levels in PKC versus PK cell lines (Figs. S2F, S2G). Taken together, these results indicate that cell lines 193 and 470 represent reasonable models for further study.

To define the role of Ccnc in the autophagic response in pancreatic cells, the appearance of autophagic protein LC3B puncta, a marker of active autophagy, were determined in 193 and 470 cell lines by IHC. The 193 and 470 cell lines were treated with the drug chloroquine (CQ), a potent stimulator of the ALP (Kang and Tang 2012). Following CQ treatment, LC3B puncta were observed in the 470 cell line but not in 193 *Ccnc^-/-^* cells (Fig. 4A). To directly monitor ALP activity, we next used Western blot analysis to distinguish between the unmodified (LC3BI) and lipidated induced LC3B species (LC3BII). Under uninduced conditions, LC3BI and LC3BII levels were similar in both cell lines (Fig. 4B, left two lanes). LC3BII levels normally increase following treatment by CQ and/or the mTOR inhibitor Torin 1 (Thoreen et al. 2009). Elevated LC3BII levels were observed in the 470 *Ccnc^+/+^* cell line following CQ treatment (Fig. 4B). However, no increase in L3CBII was observed in the 193 *Ccnc^-/-^*cells treated with either drug or in combination. These results indicate that the 193 cell line is defective for autophagy induction. To confirm that the reduced autophagic response was due to reduced Cdk8 and/or Cdk19 activity, 470 *Ccnc^+/+^*cells were treated with the Cdk8/Cdk19 inhibitor Senexin A (Galbraith et al. 2017). Treating 470 cells with Senexin A reduced LC3BII accumulation following CQ and Torin-1 treatment similar to the *Ccnc^-/-^* 193 cell line (Fig. 4C).

**Figure 4.**
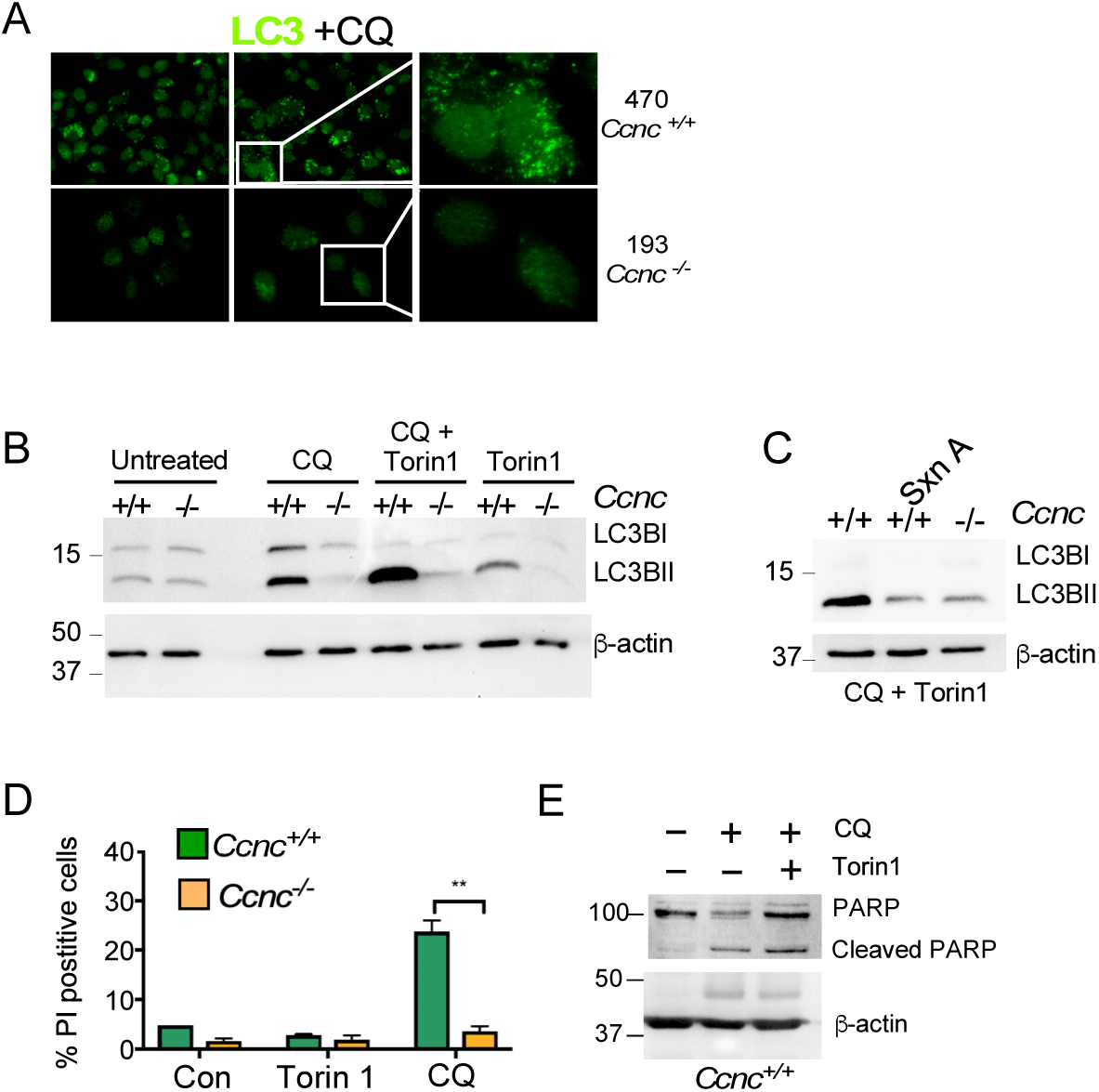
Ccnc is required for autophagic induction in pancreatic cell line. (A) IHC analysis for LC3B puncta in the *Kras^G12D/+^*;*Ccnc^+/+^* (470) and *Kras^G12D/+^*;*Ccnc^-/-^* (193) 193 cell lines following chloroquine (CQ, 50 µM, 24 h) treatment. Zoomed panels are the right. (B) Western blot analysis of the unmodified (LC3BI) and lipodated (LC3BII) species in whole cell extracts prepared from 470 and 193 cell lines with the indicated treatments of autophagy-inducing CQ (50 µM, 24 h) and/or Torin 1 (250 nM, 24 h). ß-actin served as a loading control. Molecular weight markers are indicated on the left (kDa). (C) Western blot analysis probing for LC3BI and LC3BII in extracts prepared from CQ/Torin 1 treated 470and 193 cell lines exposed to the Cdk8/Cdk19 inhibitor Senexin A as indicated. (D) Viability studies for 470 and 193 cell lines treated with chloroquine (CQ, 50 µM, 24 h) or Torin 1 (250 nM, 24 h). Non-viable, PI permeable cells were counted by fluorescent cell analysis. Percent of the population PI positive (±SD) is presented. N = 3 independent cultures. (E) Western blot probing for cleaved PARP in extracts prepared from line 470 cell line treated with autophagic inhibitors indicated. ß-actin served as a loading control. Molecular weight markers are on the left (kDa).

Finally, we determined whether either the 193 or 470 cell lines were sensitive to an autophagy inhibitor. Previous studies revealed that elevated autophagy observed in PDACs makes these cells sensitive to inhibitors (Yang et al. 2011; Kang and Tang 2012). Both cell lines were treated with either Torin 1 or CQ, and their viability measured using fluorescence cell analysis of PI uptake by dead cells. These experiments revealed no change in viability in either Torin 1 treated cell line (Fig. 4D). However, the *Ccnc^+/+^* 470 cells exhibited a significant increase in cell death following CQ treatment while the 193 cells did not. The resistance of 193 cells to CQ is consistent with our finding that these cells do not induce ALP. To further assess the nature of the cell death, PARP cleavage was measured in 470 cells treated with CQ or CQ + Torin 1. These experiments revealed enhanced PARP cleavage indicating that the cells were dying via apoptotic mechanisms. Taken together, these results confirm the role of Ccnc in autophagy gene expression and execution of the process. As autophagy is required for maintaining normal acinar homeostasis, these findings indicate that pancreatic damage in *Ccnc^PancΔ^* animals is caused, at least in part, by reduced autophagy (see Discussion).

### The 193 cell line exhibits elevated endogenous ROS and colony forming ability

Several studies found that autophagy reduces ROS production and increases genomic stability in pancreatic tissues (Kang and Tang 2012; Li et al. 2018). Depending on the concentration, ROS can serve as a signaling molecule able to enhance cell growth (Chiu and Dawes 2012) or induce damage and even cell death (Kim 2008; Klaunig 2018). Oxidation increases dihydroethidium (DHE) fluorescence permitting total cellular ROS to be monitored. We examined DHE fluorescence in immortal MEF, *Kras^G12D/+^*(470), and *Kras^G12D/+^;Ccnc^-/-^* (193) cell lines. As expected, the immortal MEF line exhibited reduced DHE oxidation compared to the transformed *Kras^G12D/+^* 470 cells (Fig. 5A). Interestingly, the *Kras^G12D/+^;Ccnc^-/-^*193 cells exhibited even higher DHE oxidation than the 470 line. We next examined mitochondrial ROS levels in these cell lines using the MitoSox oxidant sensitive dye. Consistent with the DHE results, we found elevated ROS in the mitochondrial compartment in 193 cells compared to the 470 cell line (Fig. 5B). Taken together, these results indicate that the *Kras^G12D+^;Ccnc^-/-^*193 cells are carrying a higher ROS burden than *Kras^G12D+^* cells alone.

**Figure 5.**
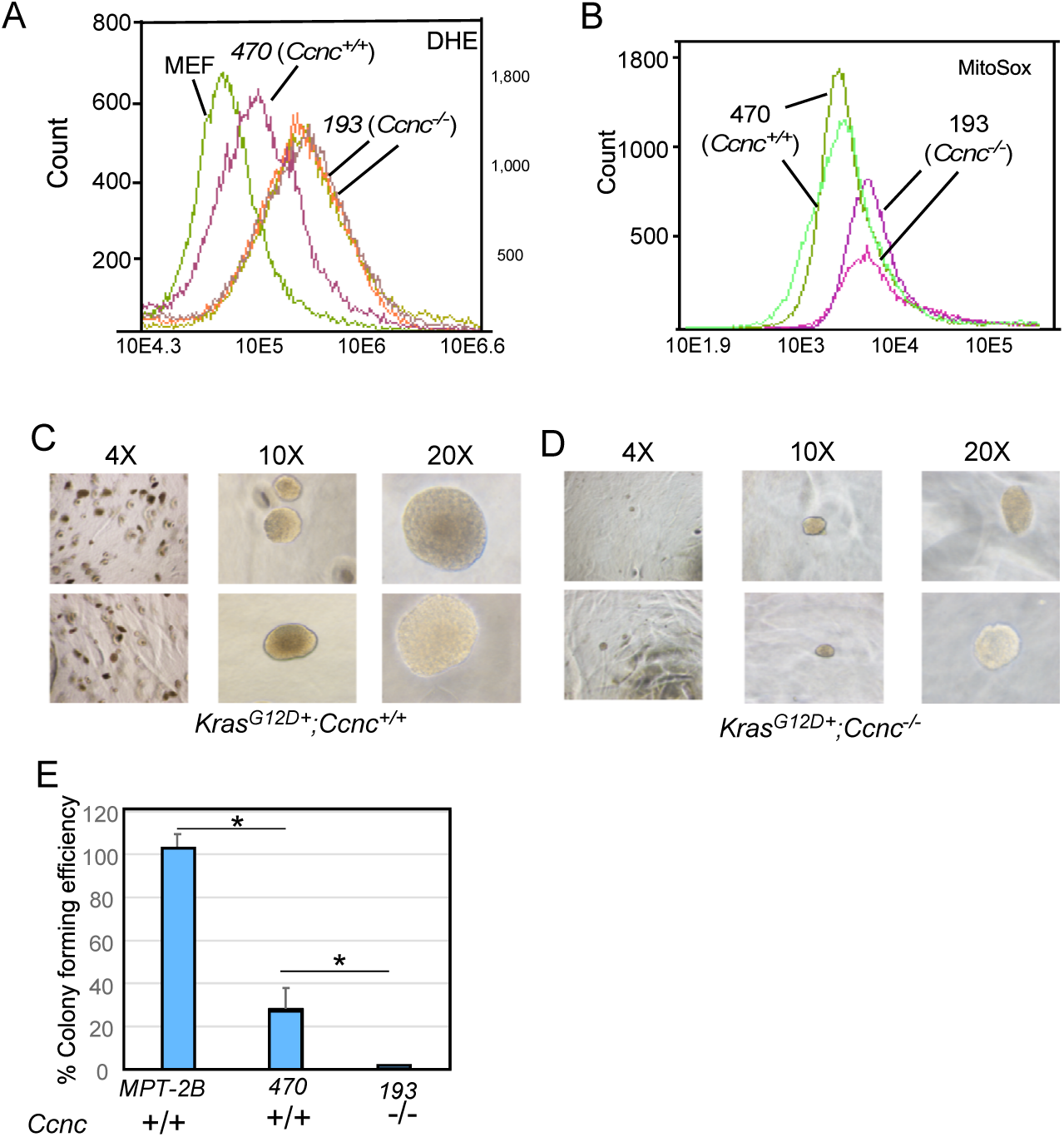
Transformed phenotypes of *Kras^G12D/+^*;*Ccnc^-/-^* cell lines. (A) Endogenous ROS levels were determined by DHE staining followed by fluorescence cell analysis. WT immortalized MEF cells are used as a control for non-transformed cell type. *Kras^G12D/+^*;*Ccnc^+/+^*(470) and *Kras^G12D/+^*;*Ccnc^-/-^* (193) traces are indicated. Biological replicates for the 193 cell line are shown. (B) Two biological replicates of the 470 and 193 cell lines were stained with MitoSox and analyzed as described in (A). (C) The MPT-2B (*Kras^G12D/+^*;*Trp53^+/-^*;*Ccnc^+/+^*) and (D) 193 (*Kras^G12D/+^*;*Ccnc^-/-^*) cells were plated in soft agar and colony forming ability imaged. Representative images following three weeks in culture are shown at the indicated magnification. (E) Colony forming efficiency of MPT-2B was set at 100% for comparing 470 and 193 cell lines. Asterisks indicate p values of < 0.05.

We next addressed whether the 193 and 470 cell lines were able to form colonies in soft agar, a common trait of transformed cells. As a positive control, we tested the colony forming ability of the MPT-2B (*Kras^G12D+^;Trp53^+/-^;Ccnc^+/+^*), a mouse cell line that demonstrated tumor forming ability following orthotopic injections. As expected from the aggressive nature of this tumor, the MPT-2B readily produced large colonies in soft agar, which was used as a standard for colony formation (100%). The 470 cell line exhibited strong colony formation (Fig. 5C, quantified in Fig. 5E) but not to the extent of the MPT-2B cells. The 193 cell line produced relatively small colonies (Fig. 5D) at lower frequency. Furthermore, no evidence of pancreatic tumor development was observed eight months after mice were injected orthotopically with the 193 cells (data not shown). Therefore, despite rapid development of aggressive PanIN lesions in PKC mice, a cell line derived from these mice exhibit a weak transformed phenotype. This may be due to the relatively early time (15 weeks) of tissue harvesting precluding the accumulation of additional mutations. Alternatively, the reduced autophagy in this cell line may retard more advance cancer stages as suggested previously (Kang and Tang 2012) (see Discussion).

### Ccnc is required for normal 26S proteasome function

The requirement of Ccnc for the ALP raised the question about how this impacted UPS activity. Reduced UPS activity has been shown to upregulate the ALP to maintain proteostasis (reviewed in (Dikic 2017)). However, studies have both supported (Wang et al. 2013) or refuted (Qiao and Zhang 2009) that autophagy inhibition stimulates UPS activity. To determine if Ccnc loss alters UPS activity in our pancreatic cell lines, extracts prepared from 193 and 470 cells were incubated with a model substrate (LLVY-AMC) that fluoresces when cleaved by the proteasome. These studies revealed that the relative velocity of the proteasome activity was 113 RFU/min for 470 cells (Fig. 6A). Interestingly, *Ccnc^-/-^*193 cell extracts exhibited significantly reduced proteasome activity (62 RFU/min). Next, we determined whether *Ccnc^-/-^* cells were still sensitive to MG-132 inhibition. In these experiments, we treated cells with the same concentration of MG-132 used to induce autophagy. MG-132 treatment reduced UPS activity in both the *Ccnc^+/+^* (64 RFU/min) and *Ccnc^-/-^* (27 RFU/min) cell lines (Fig. 6B). These results indicated that Ccnc is also required for normal proteasome function as well as ALP.

**Figure 6.**
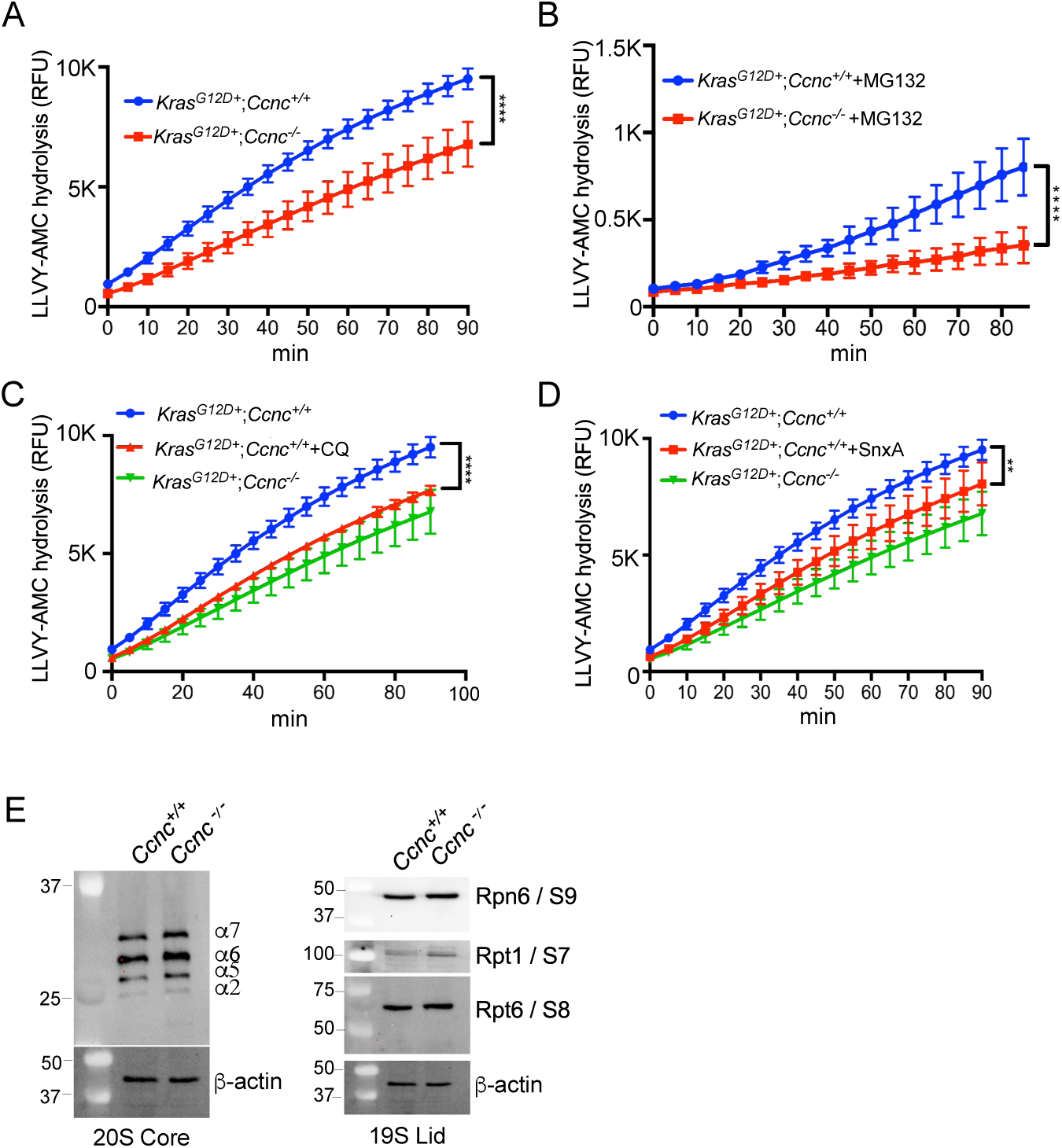
Ccnc maintains normal UBS function. (A) LLVY-AMC cleavage assays were performed with extracts prepared from 193 (*Ccnc^-/-^*) and 470 (*Ccnc^+/+^*) cell lines. The indicated cell lines were treated with the proteasome inhibitor MG132 (B), CQ (C) and Senexin A (D). The control curves for panels C and D are transferred from the experiment in panel A. (E) Western blot analysis of the indicated protein components of the 20S and 19S complexes. Molecular weight markers (kDa) are indicated on the left. In all panels, *=p<0.05; **=p<0.01; ***=p<0.005; ****=p<0.001.

One possible explanation of our results is that reduced autophagy gene expression has a negative impact on proteasome function in this background. If correct, two predictions should be true. First, inhibiting autophagy in *Ccnc^+/+^* cells should phenocopy the *Ccnc^-/-^* results. Therefore, proteasome activity was measured in 470 *Ccnc^+/+^* cells treated with CQ. These studies revealed a similar reduction in proteasome activity as observed in *Ccnc^-/-^* cells (Fig. 6C). Second, inhibiting Cdk8/Cdk19 activity with the drug Senexin A should also reduce proteasome function in *Ccnc^+/+^*cells. Senexin A treatment caused a significant reduction in proteasome activity although not to the extent observed for the *Ccnc^-/-^* cell line (Fig. 6D). These results are consistent with the model that Ccnc is required for autophagy gene transcription which reduces both autophagy and proteasome activity. Alternatively, Ccnc may directly support UPS activity by mediating transcription of proteasome component genes. To address this question, we monitored the levels of several components of the 20S core particle and the 19S regulatory lid in the 193 and 470 cell lines. These experiments did not reveal any significant difference in the selected proteins from either complex (Fig. 6E), although this does not rule out other proteasome components being down regulated. Taken together, these results reveal a new role for CKM transcriptional control in supporting both ALP and UPS activity although the latter role may be indirect.

### Reducing CKM activity sensitizes tumor cell lines to proteasome inhibitors

Proteotoxic stress stimulates both the UPS and ALP systems (Dai et al. 2007; Amaravadi et al. 2016). Therefore, we tested the sensitivity of the *Ccnc^-/-^*cell line to the UPS inhibitors bortezomib (BZ) or MG-132 utilizing Annexin V staining to monitor iRCD. BZ or MG-132 treatment of the 470 (*Ccnc^+/+^*) cell lines resulted in three and five-fold increase in cell death versus untreated control cells (Fig. 7A). Repeating these experiments with the 193 cell line revealed an increased sensitivity to BZ or MG-132 treatment compared to 470 cells. These results indicate that loss of Ccnc activity sensitizes cells to proteasome inhibitors. To determine if this hypersensitivity was expressed in other tumor types, *CCNC* was deleted via CRISPR protocols in HeLa (cervical) and HCT116 (colon) human cancer cell lines (Fig. S4). *CCNC^-/-^* HeLa cells exhibited a threefold increase in cell death following BZ treatment compared to the *CCNC^+/+^* parental cells (Fig. 7B). A similar, but less dramatic effect, was observed in BZ treated HCT116 *CCNC^-/-^* cells (Fig. 7C). Taken together, these studies indicate that inactivating CCNC results in a general hypersensitivity to proteasome inhibitors.

**Figure 7.**
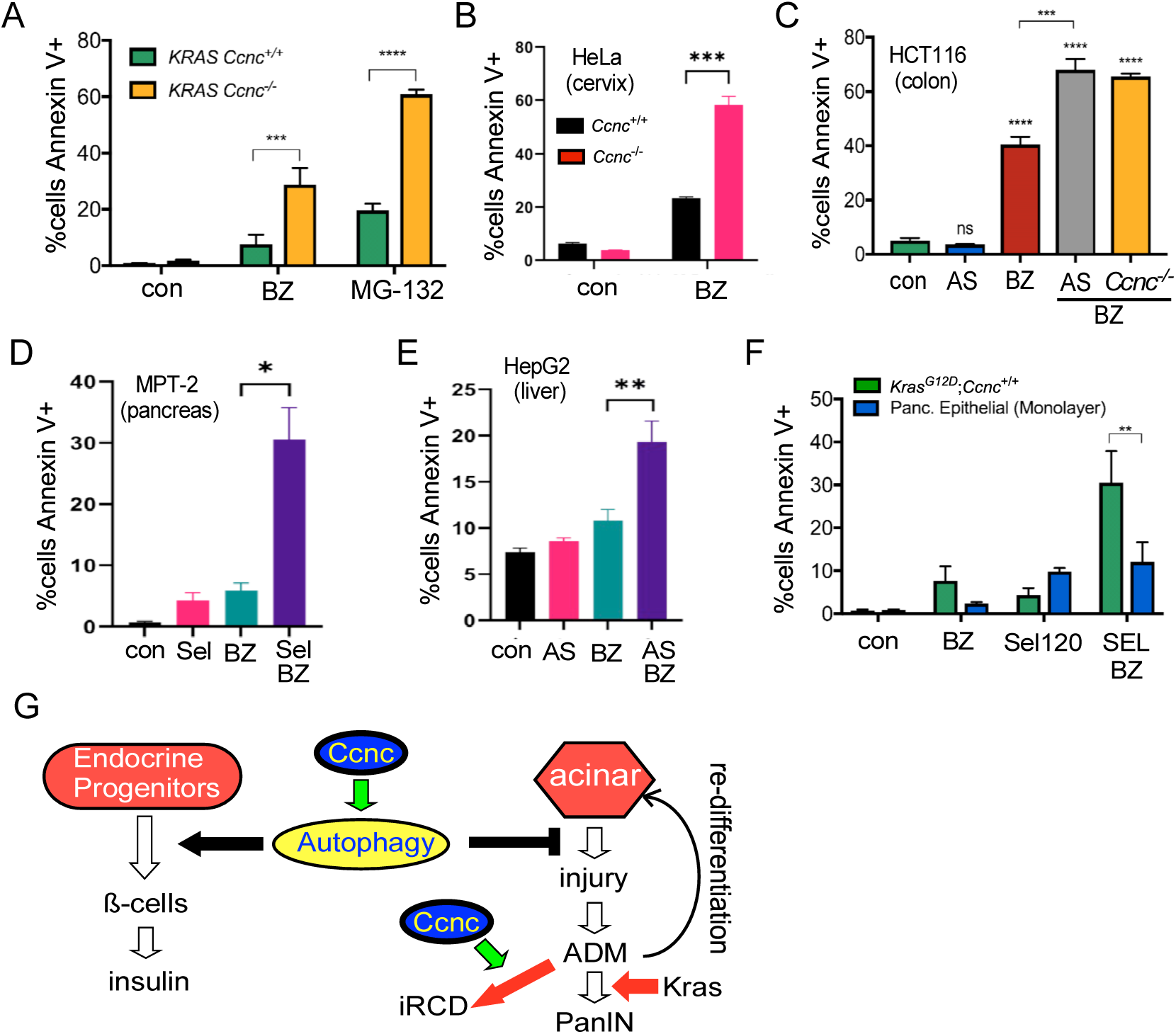
Reduced CKM function enhances sensitivity to proteasome inhibitors. (A) 470 (*Kras^G12D/+^*;*Ccnc^+/+^*) and 193 (*Kras^G12D/+^*;*Ccnc^-/-^*) cells were treated with the proteasome inhibitors bortezomib (BZ, 100 nM) or MG-132 (250 nM) and percent of the population Annexin V positive;PI negative determined. (B) *Ccnc^+/+^*or *Ccnc^-/-^* HeLa cells were treated with BZ and analyzed as before. (C) *Ccnc^+/+^* and *Ccnc^-/-^* HCT-116 cells were treated with the CDK8/19 inhibitor AS-2863619 (AS, 1 µM) and/or BZ as indicated. HCT-116*^Ccnc-/-^* cells were treated with only BZ. (D) MPT-2 cells were treated with the CDK8/19 inhibitor Sel-120 (Sel, 1 µM) and/or BZ. (E) HepG2 cells were treated as described in (C). (F) MPT-2B or non-transformed pancreatic epithelial cells were treated as indicated. In all panels, *=p<0.05; **=p<0.01; ***=p<0.005; ****=p<0.001. (G) Autophagy supports ß-cell differentiation, islet formation and insulin production thus protecting the animal from hyperglycemia and shortened animal life. In addition, autophagy protects acinar cells from proteotoxic stress by removing misfolded proteins and aggregates. ALP dysfunction induces ADM dedifferentiation where it can redifferentiation back to the acinar lineage. The second role is related to the ability of activated Kras^G12D^ to drive proliferating ADM cells toward PanIN formation. This process can alternatively lead to Ccnc-dependent cell death thus blunting disease progression.

We next tested whether CDK8/CDK19 inhibitors phenocopied the *CCNC^-/-^* allele with respect to proteasome inhibitor hypersensitivity. First, HCT-116 *CCNC^+/+^* cells were treated for 24 h with the CDK8/CDK19 inhibitor AS-2863619 (AS) (Akamatsu et al. 2019) prior to BZ addition. As a control, AS treatment alone had no significant impact on HCT-116 viability (Fig. 7C). However, BZ addition increased AS treated HCT-116 cell death to levels observed for the *CCNC^-/-^* derivative (Fig. 7C). Finally, these studies were repeated with two additional cell lines, MPT-2B and the liver cancer cell line HepG2. For the MPT-2B cells, a different CDK8/CDK19 inhibitor (Sel-120 (Rzymski et al. 2017), was used. In both experiments, inhibiting Cdk8/Cdk19 activity increased BZ sensitivity (Fig. 7D and 7E). Importantly, treating SV-40 immortalized normal pancreatic epithelial cells with this regimen did not significantly increase toxicity over proteasome inhibitor alone (Fig. 7F). These findings point to a general enhanced sensitivity of this approach to transformed versus non-transformed cells. The analysis of additional cell types revealed that a thyroid cell line (D445) or normal human lung fibroblasts (IMR90) cells were modestly sensitized to BZ following treatment with the Cdk8/Cdk19 inhibitor (Fig. S5A and S5B). Finally, we found that the 193 cell line was more sensitive than 470 cells to BTZ over multiple drug concentrations (Fig. S5C). Importantly, this study also revealed a several fold elevated sensitivity to BZ by the 193 cells compared to proliferating normal pancreatic epithelial cells. This difference was more striking when quiescent pancreatic epithelial cells were compared to the 193 cell line. Taken together, these results indicate that inhibiting CDK9/CDK19 represents a reliable strategy to specifically induce tumor cell hypersensitivity to proteasome inhibitors.

## Discussion

This report identifies a role for Ccnc in supporting both basal and induced autophagic gene transcription. We find that deleting *Ccnc* results in several phenotypes previously observed in autophagy pancreatic ablation models including defective islet development, reduced insulin production and ADM. The presence of oncogenic Kras (Kras^G12D^) resulted in the accelerated appearance of extensive ADM and PanIN lesions indicating that Ccnc prevents early events in pancreatic cancer initiation. Finally, loss of Ccnc activity results in a reduction in both ALP and UPS activity rendering cells hypersensitive to proteasome inhibitors perhaps providing a new avenue to attack pancreatic cancers. These findings indicate that Ccnc plays a pivotal role in both pancreatic function and disease suppression.

The present study revealed two roles for Ccnc in the pancreas (Fig.7G). First, autophagy is required for normal ß-cell islet development (Jung et al. 2008; Marasco and Linnemann 2018) and preventing acinar cell damage (Antonucci et al. 2015; Diakopoulos et al. 2015; Habtezion et al. 2019). We found that *Ccnc* is required for normal islet development, insulin production and preventing ADM formation. A previous study found that p53 induced the transcription of the macroautophagy gene *DRAM1* (Crighton et al. 2006). Similarly, we found that Ccnc is required for both steady state and induced *Dram1* transcription. As Ccnc-Cdk8 has previously been identified as a co-activator with p53 for inducing p21 transcription (Donner et al. 2007), this may indicate that a similar regulatory mechanism is present at *DRAM1* and other autophagy gene promoters.

The role CKM components play in murine development is complicated. For example, deleting *Cdk8* results in embryo arrest at the preimplantation stage (Westerling et al. 2007). Conversely, *Cdk19^-/-^* mice exhibit a normal lifespan (Dickinson et al. 2017). In the middle of these extremes, *Ccnc^-/-^* mice arrest at day 9.5 at the onset of organogenesis ((Li et al. 2014) and our unpublished results). However, pancreatic *Cdk8* ablation in combination with *Kras^G12D^* expression did not increase the presence of pre-cancerous lesions above Kras^G12D^ expression alone. This result was somewhat surprising but suggests that either Cdk8 activity is replaced by Cdk19 (Fant and Taatjes 2019; Chen et al. 2023). However, as *Cdk19^-/-^* animals develop normally, its requirement alone seems unlikely. Therefore, deleting both kinases would most likely be required to phenocopy *Ccnc^-/-^*cells.

However, the phenotypes associated with loss of Ccnc function were more severe than deleting *Atg5* or *Atg7*. For example, *Atg5* or *Atg7* pancreatic ablation led to shortened lifespan (median ∼16 weeks) although 40% of the animals exhibited a normal life span (Rosenfeldt et al. 2013). Animals harboring *Ccnc* pancreatic deletion did not survive past 15 weeks. In addition, PKC animals exhibited more accelerated rate of ADM and PanIN lesions compared to pancreata harboring *Kras^G12D^* and *Atg5^-/-^* alleles. These results suggested that Ccnc plays a role(s) in pancreatic biology in addition to maintaining autophagy. This second role may be related to the mitochondrial function of Ccnc contributing to restricting neoplastic growth (Fig. 7G). Activated Kras increases mitochondrial derived ROS through elevated metabolism associated with transformed growth (Liou et al. 2016). In many cancers, ROS are maintained at levels that enhance mutagenesis and proliferation but below that which induces iRCD (Klaunig 2018). Our previous studies revealed that *Ccnc* deletion inhibits ROS-induced iRCD (Jezek et al. 2019). Therefore, loss of Ccnc activity could allow continued growth of Kras^G12D^-driven cell division even in the presence of normally toxic ROS levels. Consistent with this possibility, we find that the *Kras^G12D^*, *Ccnc^-/-^* cell line exhibits higher endogenous and mitochondrial ROS concentrations than a *Kras^G12D^* tumor cell line.

A connection between hyperglycemia and cancer has been proposed in that tumor growth is stimulated by excessive serum glucose (Garcia-Jimenez et al. 2014; Cosmin Stan and Paul 2024). Consistent with this possibility, mice harboring pancreatic tumors deleted for the autophagy gene *Kras^G12D/+^*;*Atg7^-/-^;Tp53^-/-^* exhibited increased glucose uptake resulting in elevated glycolysis to support anabolic pathways (Rosenfeldt et al. 2013). Interestingly, PC animals exhibit elevated serum glucose, but PKC mice do not despite exhibiting poorly developed islets. A possible explanation for these findings is that the presence of extensive PanIN lesions in PKC mice also metabolizes glucose at an elevated rate thus reducing serum glucose levels. This would suggest that the metabolic reprogramming associated with PDAC may be manifested earlier in precancerous lesions lacking Ccnc.

PDACs are usually discovered following metastasis making surgical options difficult thus elevating the reliance on chemotherapeutic approaches. However, the success rates of current regimens are disappointing. Therefore, new approaches are necessary to improve patient outcome. Due to the unregulated growth associated with transformation, elevated concentrations of damaged or misfolded proteins generating proteotoxic stress stimulates both UPS and ALP activity (Dai et al. 2007; Amaravadi et al. 2016). Increased proteostasis is important for cancer cells survival as *KRAS* transformed cells become addicted to this activity with cells becoming hypersensitive to proteasome inhibition (Luo et al. 2009). Although proteasome inhibitors have proven useful in some hematopoietic malignancies, they have been less successful in solid tumors including pancreatic cancers (reviewed in (Murugan and Voutsadakis 2022)). One possible explanation for this result is that solid tumors require an additional sensitization step. Our finding that the activity of both the ALP and UPS are reduced following *Ccnc* ablation or Cdk8/Cdk19 inhibition may sensitize tumors to proteasome inhibitors. Therefore, our results suggest that even if *CCNC* deletion is not directly involved in tumorigenesis, its loss may still be useful for targeting tumors with proteasome inhibitors.

## MATERIALS AND METHODS

### Animals

All animal experiments were conducted in accordance with institutional animal care and use committee (IACUC) review at Fox Chase Cancer Center (FCCC) and RowanSOM. The mice used in this study were backcrossed at least ten generations into the C57BL/6 background. The Cdk8 floxed mouse line *Cdk8^tm1a(EUCOMM)Hmgu^*was from the Mouse Clinical Institute as part of the Constitutive Knockout/Conditional Knockout (KO-cKO pc) project at the Institut Clinique de la Souris. This mouse was first mated with a strain harboring the FLP recombinase under the control of the Rosa promoter [B6.129S4-Gt(ROSA)26Sortm2(FLP*)Sor/J] to remove the selection marker (neomycin resistance) from the Cdk8 floxed allele. Expression of the cre recombinase deleted exon5, resulting in a frameshift mutation. The *Pdx1-Cre;LSL-Kras^G12D^* strain was originally obtained from Dr. Owen Sansom (Beatson Institute for Cancer Research, Glasgow). The floxed *Ccnc* (*Ccnc^fl/fl^*) strain was described previously (Wang et al. 2015). Littermates with the indicated genotypes were used as controls.

### Cell culture

MPT-2B and 470 cell lines were cultured at 5% CO_2_ in DMEM supplemented with 10% FBS. The 193 cell line was cultured in low calcium medium (DMEM/F-12 1:1 [Gibco, special order medium with no calcium] supplemented with 5% chelated horse serum, 20 ng/ml EGF, 100 ng/ml cholera toxin, 10 µg/ml insulin, 0.5 µg/ml hydrocortisone, 0.04 mM calcium chloride, dihydrous, 100 U/ml penicillin, 100 µg/ml streptomycin, 10 µg/ml ciprofloxacin, and 0.25 µg/ml amphotericin B). These three cell lines were generated from pancreata of mice with the indicated genotypes in the FCCC Cell Culture Facility.

### Mouse genotyping

The floxed *Ccnc* (*Ccnc^fl^*) (Wang et al. 2015), *Kras^G12D^* and the pancreas-specific Cre (*Pdx1*-Cre) alleles have been previously described (Hingorani et al. 2003; Hingorani et al. 2005). Genotyping at Rowan University was accomplished using genomic tail DNA purified using the Phire genotyping kit and a split PCR method was run for 35 cycles (Thermo Scientific, Rockford, IL, USA). Genotyping at FCCC was performed by the Mouse Genotyping Facility from genomic tail DNA using Promega GoTaq Green Master Mix or Sigma JumpStart Taq ReadyMix. The genotyping primers are listed in the Key Resources Table S1.

### Immunohistochemistry

H&E staining was performed as previously described (Antico Arciuch et al. 2011). Ki-67 staining was conducted essentially as previously described (Saad et al. 2006). Immediately after euthanizing the mice, pancreata were collected and all connective tissue was removed. The tissues were then washed twice in formalin and fixed in formalin overnight. Then the tissues were stored in 70% ethanol for future examination. All the tissues were embedded in paraffin and sectioned at 6 µm. Sections were subjected to antigen retrieval in 0.1 mM sodium citrate and counterstained with hematoxylin. All stained sections were photographed at 40-200x magnifications and analyzed using ImageJ software.

### Western blot analysis

Mouse cells were homogenized in RIPA buffer (150 mM NaCl, 50 mM Tris, 1% Nonidet P-40 substitute, 0.5% sodium deoxycholate, 0.1% SDS, pH 8) containing 1% protease inhibitor cocktail (PIC) (Sigma P8340, St. Louis, MO USA), 1% EZBlock phosphatase inhibitor cocktail IV (BioVision, Milpitas, CA USA), 10 mM NaF, 10 mM β-glycerol phosphate, and 2 mM Na_3_VO_4_. Homogenates were incubated for 2 h at 4 °C and centrifuged at 14,000 × g for 20 min at 4 °C to separate soluble proteins from aggregates and cell debris. Protein concentrations were determined by Bradford assay (Bio-Rad, Hercules, CA USA). Samples were dissolved in sample buffer (SB) (100 mM Tris, 4% SDS, 20% glycerol, 2 mg/ml bromophenol blue, pH 6.8) supplemented with 100 mM DTT, boiled for 5 min, and separated by SDS-PAGE then probed with the indicated antibodies. Western blot signals were visualized using CDP-STAR (CDP*) substrate and signals were visualized using an iBright FL1500 Imaging System (Thermo).

### RT-qPCR analysis

Total RNA was prepared from the cell samples using Monarch® Total RNA Miniprep Kit. On-column DNase treatment was performed to eliminate contaminating DNA during RNA extraction. Total RNA (500 ng) was converted to cDNA using the ThermoFisher™ Maxima cDNA Synthesis kit. The cDNA from each sample (1/100 dilution) was subjected to qPCR amplification using ThermoFisher™ PowerSYBR™ Green PCR Master Mix and a StepOne™ Real Time PCR System. These assays were conducted with three independent preparations assayed in duplicate. *Gapdh* was used as the internal standard for comparative (ΔΔC_T_) quantitation (Livak and Schmittgen 2001). Statistical significance was determined via Student’s *t*-test analysis. P values <0.05 were considered significant. Primer sequences for RT-qPCR analysis can be found in Table S1.

### CRISPR CCNC knockout protocol

*CCNC* was depleted in the HeLa and HCT116 cell lines by dually transfecting CRISPR Cas12a plasmids (pTE4398, Addgene) containing gRNA directed to Exon 1 and Exon 2 (pSW496 and pSW497, respectively). Transfections were performed by electroporation using the Neon Electroporator (ThermoFisher) as described by the manufacturer. Briefly, trypsin/EDTA treated cells harvested at 50% confluent were washed then resuspended at 1x10^6^ cells/mL in R buffer. Cells were electroporated in 10 µL tip at 1250 V with 20 ms pulse width. The transfectants were selected for 7 days with puromycin and then clonally diluted into 96 well plates. Individual clones were then analyzed by Western blot for the presence of CCNC.

### Soft agar assays

The cell lines indicated in tissue culture medium were resuspended in 1.5 ml of 0.3% noble agar in Hank’s buffer (Millapore-Sigma H6648) at 37°. The cell suspensions were immediately plated into a 24-well plate with 0.5 ml of 0.6% bottom agar medium already solidified. Once the suspensions solidified (30 min), the plates were incubated at 37° for three weeks. 100 µL of medium was added to each well as necessary. The cells were fixed with methanol and stained with 0.15% crystal violet prior to imaging. The percentage of the cells plated forming macroscopic colonies was calculated.

### Plasma glucose analysis

Plasma was obtained at 2-week intervals from mice starting at 4 weeks of age by retroorbital bleed into heparinized tubes. Plasma was cryopreserved at -80°C and glucose concentrations were performed at 1:100 dilution using a Colorimetric Glucose Assay Kit from Cell Biolabs, Inc. (#STA-680; San Diego, CA) according to manufacturer’s instructions.

### Proteasome activity assays

26S proteasome activity was determined in adherent cells grown to ∼75% confluency and lysed using buffer A (25 mM Tris, pH 7.4, 10 mM MgCl2, 10% glycerol) supplemented with fresh 1 mM ATP, 1 mM DTT, and 5 μL (for 500 μL of buffer) of protease inhibitor and phosphatase inhibitor. Cells were centrifuged, resuspended in 75 μL of lysis buffer A, and incubated for 30 min at 4⁰ with gentle shaking. Cell debris was removed by centrifugation (15K x g) for 4 min at 4⁰ and the resulting lysates were retained. Each assay contained 25 µg of protein as determined by Bradford protein assays. To monitor proteasomal-independent fluorescence, the proteasome inhibitor MG-132 (72.5 μM) was added to each control well and non-specific cleavage of the substrate subtracted from the experimental samples. Finally, the substrate buffer (lysis buffer A, 200 μM Suc LLVY, 1 mM DTT, and 1 mM ATP) was added to each well. Fluorescence was immediately monitored via a plate reader. Fluorescent readings (excitation = 360, emission = 400) were taken every 5 minutes for 90 minutes at 37⁰ with gentle agitation. All assays were performed with three biological repeates with two technical replicates. In these assays, Senexin A (0.4 mM) and chloroquine (100 µM) was added as indicated 24 h prior to cell harvesting.

### Statistical analysis

The student’s *t* test was applied to determine statistical differences for the data presented. P values <0.05 were considered significantly different in these assays. The Kaplan-Meier survival curves were calculated using GraphPad. Actual calculations of p values are provided in each figure as denoted in the legend.

## Abbreviations

Cdk8: cyclin dependent kinase 8
PDAC: pancreatic ductal adenocarcinoma
H&E: hematoxylin and eosin
ROS: reactive oxygen species
PanIN: pancreatic intraepithelial neoplasias
ADM: acinar-to-ductal metaplasia
iRCD: intrinsic regulated cell death
MEF: Mouse embryonic fibroblasts
ALP: autophagy lysosome pathway
UPS: ubiquitin-proteasome system.

## Acknowledgements (Research support)

## Acknowledgements

The authors appreciate expert technical support from the FCCC Histopathology, Cell Culture, Laboratory Animal, Bioinformatics & Biostatistics, and Mouse Genotyping Facilities, specifically the work of Catherine (Cass) Renner, Jirong (Jenny) Zhang, Fang Jin, Suraj Peri, Irina Shchaveleva, Justin Rambert, Tyanah Yisrael, Lauren Heisey, Mark Tague, Simon Tarpinian, and Rita Michielli. We also thank Dr. Jonathan Chernoff for the SV40-importalized mouse pancreatic epithelial cell line and Dr. Edna Cukierman for the normal pancreatic cell line. We thank Drs. Edna. Cuckierman, Igor Astsaturov and members of the Marvin and Concetta Greenberg Pancreatic Cancer Institute at FCCC for advice.

## Acknowledgements (Funding)

This work was supported by grants from the National Institutes of Health awarded to R.S. (GM113052) and the FCCC Comprehensive Cancer Center Support Grant (CA06927) in support of the Histopathology, Cell Culture, Laboratory Animal and the Mouse Genotyping Facilities at FCCC. Additional support was provided by the Boye Foundation and the New Jersey Health Foundation (to R.S.) and from the Martin and Concetta Greenberg Pancreatic Cancer Institute at FCCC and Pennsylvania DOH Health Research Formula Funds (to K.S.C.).

## CRediT author statement

Sara Hanley, Investigation, Formal analysis; Kathy Q. Cai, Investigation; Formal analysis; Stephen D. Willis, Investigation, Formal analysis; David C. Stieg, Investigation, Formal analysis; Andres J. Klein-Szanto, Writing-Review and Editing; Kerry S. Campbell, Writing, Review and Editing, Supervision, Funding acquisition, Resources; Randy Strich, Writing, Original Draft, Review and Editing, Supervision, Funding acquisition, Resources.

## References

Akamatsu M, Mikami N, Ohkura N, Kawakami R, Kitagawa Y, Sugimoto A, Hirota K, Nakamura N, Ujihara S, Kurosaki T et al. 2019. Conversion of antigen-specific effector/memory T cells into Foxp3-expressing T(reg) cells by inhibition of CDK8/19. Sci Immunol 4.

Amaravadi R, Kimmelman AC, White E. 2016. Recent insights into the function of autophagy in cancer. Genes Dev 30: 1913–1930.

Antico Arciuch VG, Russo MA, Dima M, Kang KS, Dasrath F, Liao XH, Refetoff S, Montagna C, Di Cristofano A. 2011. Thyrocyte-specific inactivation of p53 and Pten results in anaplastic thyroid carcinomas faithfully recapitulating human tumors. Oncotarget 2: 1109–1126.

Antonucci L, Fagman JB, Kim JY, Todoric J, Gukovsky I, Mackey M, Ellisman MH, Karin M. 2015. Basal autophagy maintains pancreatic acinar cell homeostasis and protein synthesis and prevents ER stress. Proc Natl Acad Sci U S A 112: E6166–6174.

Barette C, Jariel-Encontre I, Piechaczyk M, Piette J. 2001. Human cyclin C protein is stabilized by its associated kinase cdk8, independently of its catalytic activity. Oncogene 20: 551–562.

Bourbon HM. 2008. Comparative genomics supports a deep evolutionary origin for the large, four-module transcriptional mediator complex. Nucleic Acids Res 36: 3993–4008.

Chen M, Li J, Zhang L, Wang L, Cheng C, Ji H, Altilia S, Ding X, Cai G, Altomare D et al. 2023. CDK8 and CDK19: positive regulators of signal-induced transcription and negative regulators of Mediator complex proteins. Nucleic Acids Res 51: 7288–7313.

Chiu J, Dawes IW. 2012. Redox control of cell proliferation. Trends Cell Biol 22: 592–601.

Choi AM, Ryter SW, Levine B. 2013. Autophagy in human health and disease. N Engl J Med 368: 651–662.

Cosmin Stan M, Paul D. 2024. Diabetes and Cancer: A Twisted Bond. Oncol Rev 18: 1354549.

Crighton D, Wilkinson S, O’Prey J, Syed N, Smith P, Harrison PR, Gasco M, Garrone O, Crook T, Ryan KM. 2006. DRAM, a p53-induced modulator of autophagy, is critical for apoptosis. Cell 126: 121–134.

Dai C, Whitesell L, Rogers AB, Lindquist S. 2007. Heat shock factor 1 is a powerful multifaceted modifier of carcinogenesis. Cell 130: 1005–1018.

Dannappel MV, Sooraj D, Loh JJ, Firestein R. 2018. Molecular and in vivo Functions of the CDK8 and CDK19 Kinase Modules. Front Cell Dev Biol 6: 171.

De La OJ, Emerson LL, Goodman JL, Froebe SC, Illum BE, Curtis AB, Murtaugh LC. 2008. Notch and Kras reprogram pancreatic acinar cells to ductal intraepithelial neoplasia. Proc Natl Acad Sci U S A 105: 18907–18912.

Diakopoulos KN, Lesina M, Wormann S, Song L, Aichler M, Schild L, Artati A, Romisch-Margl W, Wartmann T, Fischer R et al. 2015. Impaired autophagy induces chronic atrophic pancreatitis in mice via sex- and nutrition-dependent processes. Gastroenterology 148: 626–638 e617.

Dickinson ME, Flenniken AM, Ji X, Teboul L, Wong MD, White JK, Meehan TF, Weninger WJ, Westerberg H, Adissu H et al. 2017. Corrigendum: High-throughput discovery of novel developmental phenotypes. Nature 551: 398.

Dikic I. 2017. Proteasomal and Autophagic Degradation Systems. Annu Rev Biochem 86: 193–224.

Donner AJ, Szostek S, Hoover JM, Espinosa JM. 2007. CDK8 is a stimulus-specific positive coregulator of p53 target genes. Mol Cell 27: 121–133.

Fant CB, Taatjes DJ. 2019. Regulatory functions of the Mediator kinases CDK8 and CDK19. Transcription 10: 76–90.

Feng Y, He D, Yao Z, Klionsky DJ. 2014. The machinery of macroautophagy. Cell Res 24: 24–41.

Galbraith MD, Andrysik Z, Pandey A, Hoh M, Bonner EA, Hill AA, Sullivan KD, Espinosa JM. 2017. CDK8 Kinase Activity Promotes Glycolysis. Cell Rep 21: 1495–1506.

Ganesan V, Willis SD, Chang KT, Beluch S, Cooper KF, Strich R. 2019. Cyclin C directly stimulates Drp1 GTP affinity to mediate stress-induced mitochondrial hyperfission. Mol Biol Cell 30: 302–311.

Garcia-Jimenez C, Garcia-Martinez JM, Chocarro-Calvo A, De la Vieja A. 2014. A new link between diabetes and cancer: enhanced WNT/beta-catenin signaling by high glucose. J Mol Endocrinol 52: R51–66.

Grumati P, Dikic I. 2018. Ubiquitin signaling and autophagy. J Biol Chem 293: 5404–5413.

Gukovskaya AS, Gukovsky I, Algul H, Habtezion A. 2017. Autophagy, Inflammation, and Immune Dysfunction in the Pathogenesis of Pancreatitis. Gastroenterology 153: 1212–1226.

Habbe N, Shi G, Meguid RA, Fendrich V, Esni F, Chen H, Feldmann G, Stoffers DA, Konieczny SF, Leach SD, Maitra A. 2008. Spontaneous induction of murine pancreatic intraepithelial neoplasia (mPanIN) by acinar cell targeting of oncogenic Kras in adult mice. Proc Natl Acad Sci U S A 105: 18913–18918.

Habtezion A, Gukovskaya AS, Pandol SJ. 2019. Acute Pancreatitis: A Multifaceted Set of Organelle and Cellular Interactions. Gastroenterology 156: 1941–1950.

Herreros-Villanueva M, Hijona E, Cosme A, Bujanda L. 2012. Mouse models of pancreatic cancer. World J Gastroenterol 18: 1286–1294.

Hingorani SR, Petricoin EF, Maitra A, Rajapakse V, King C, Jacobetz MA, Ross S, Conrads TP, Veenstra TD, Hitt BA et al. 2003. Preinvasive and invasive ductal pancreatic cancer and its early detection in the mouse. Cancer Cell 4: 437–450.

Hingorani SR, Wang L, Multani AS, Combs C, Deramaudt TB, Hruban RH, Rustgi AK, Chang S, Tuveson DA. 2005. Trp53R172H and KrasG12D cooperate to promote chromosomal instability and widely metastatic pancreatic ductal adenocarcinoma in mice. Cancer Cell 7: 469–483.

Hruban RH, Iacobuzio-Donahue C, Wilentz RE, Goggins M, Kern SE. 2001. Molecular pathology of pancreatic cancer. Cancer J 7: 251–258.

Jezek J, Chang KT, Joshi AM, Strich R. 2019. Mitochondrial translocation of cyclin C stimulates intrinsic apoptosis through Bax recruitment. EMBO Rep 20: e47425.

Jung HS, Chung KW, Won Kim J, Kim J, Komatsu M, Tanaka K, Nguyen YH, Kang TM, Yoon KH, Kim JW et al. 2008. Loss of autophagy diminishes pancreatic beta cell mass and function with resultant hyperglycemia. Cell Metab 8: 318–324.

Kang R, Tang D. 2012. Autophagy in pancreatic cancer pathogenesis and treatment. Am J Cancer Res 2: 383–396.

Katsuragi Y, Ichimura Y, Komatsu M. 2015. p62/SQSTM1 functions as a signaling hub and an autophagy adaptor. FEBS J 282: 4672–4678.

Kim H. 2008. Cerulein pancreatitis: oxidative stress, inflammation, and apoptosis. Gut Liver 2: 74–80.

Klaunig JE. 2018. Oxidative Stress and Cancer. Curr Pharm Des 24: 4771–4778.

Kleeff J, Whitcomb DC, Shimosegawa T, Esposito I, Lerch MM, Gress T, Mayerle J, Drewes AM, Rebours V, Akisik F et al. 2017. Chronic pancreatitis. Nat Rev Dis Primers 3: 17060.

Laussmann MA, Passante E, Dussmann H, Rauen JA, Wurstle ML, Delgado ME, Devocelle M, Prehn JH, Rehm M. 2011. Proteasome inhibition can induce an autophagy-dependent apical activation of caspase-8. Cell Death Differ 18: 1584–1597.

Li N, Fassl A, Chick J, Inuzuka H, Li X, Mansour MR, Liu L, Wang H, King B, Shaik S et al. 2014. Cyclin C is a haploinsufficient tumour suppressor. Nat Cell Biol 16: 1080–1091.

Li X, Liang M, Jiang J, He R, Wang M, Guo X, Shen M, Qin R. 2018. Combined inhibition of autophagy and Nrf2 signaling augments bortezomib-induced apoptosis by increasing ROS production and ER stress in pancreatic cancer cells. Int J Biol Sci 14: 1291–1305.

Liou GY, Doppler H, DelGiorno KE, Zhang L, Leitges M, Crawford HC, Murphy MP, Storz P. 2016. Mutant KRas-Induced Mitochondrial Oxidative Stress in Acinar Cells Upregulates EGFR Signaling to Drive Formation of Pancreatic Precancerous Lesions. Cell Rep 14: 2325–2336.

Livak KJ, Schmittgen TD. 2001. Analysis of relative gene expression data using real-time quantitative PCR and the 2(-Delta Delta C(T)) Method. Methods 25: 402–408.

Luo J, Emanuele MJ, Li D, Creighton CJ, Schlabach MR, Westbrook TF, Wong KK, Elledge SJ. 2009. A genome-wide RNAi screen identifies multiple synthetic lethal interactions with the Ras oncogene. Cell 137: 835–848.

Mallen-St Clair J, Soydaner-Azeloglu R, Lee KE, Taylor L, Livanos A, Pylayeva-Gupta Y, Miller G, Margueron R, Reinberg D, Bar-Sagi D. 2012. EZH2 couples pancreatic regeneration to neoplastic progression. Genes Dev 26: 439–444.

Marasco MR, Linnemann AK. 2018. beta-Cell Autophagy in Diabetes Pathogenesis. Endocrinology 159: 2127–2141.

Morton JP, Timpson P, Karim SA, Ridgway RA, Athineos D, Doyle B, Jamieson NB, Oien KA, Lowy AM, Brunton VG et al. 2010. Mutant p53 drives metastasis and overcomes growth arrest/senescence in pancreatic cancer. Proc Natl Acad Sci U S A 107: 246–251.

Murtaugh LC, Keefe MD. 2015. Regeneration and repair of the exocrine pancreas. Annu Rev Physiol 77: 229–249.

Murugan NJ, Voutsadakis IA. 2022. Proteasome regulators in pancreatic cancer. World J Gastrointest Oncol 14: 38–54.

Qiao L, Zhang J. 2009. Inhibition of lysosomal functions reduces proteasomal activity. Neurosci Lett 456: 15–19.

Rosenfeldt MT, O’Prey J, Morton JP, Nixon C, MacKay G, Mrowinska A, Au A, Rai TS, Zheng L, Ridgway R et al. 2013. p53 status determines the role of autophagy in pancreatic tumour development. Nature 504: 296–300.

Rzymski T, Mikula M, Zylkiewicz E, Dreas A, Wiklik K, Golas A, Wojcik K, Masiejczyk M, Wrobel A, Dolata I et al. 2017. SEL120-34A is a novel CDK8 inhibitor active in AML cells with high levels of serine phosphorylation of STAT1 and STAT5 transactivation domains. Oncotarget 8: 33779–33795.

Saad AG, Kumar S, Ron E, Lubin JH, Stanek J, Bove KE, Nikiforov YE. 2006. Proliferative activity of human thyroid cells in various age groups and its correlation with the risk of thyroid cancer after radiation exposure. J Clin Endocrinol Metab 91: 2672–2677.

Siegel RL, Miller KD, Wagle NS, Jemal A. 2023. Cancer statistics, 2023. CA Cancer J Clin 73: 17–48.

Stieg DC, Chang KT, Cooper KF, Strich R. 2019. Cyclin C Regulated Oxidative Stress Responsive Transcriptome in Mus musculus Embryonic Fibroblasts. G3 (Bethesda) 9: 1901–1908.

Stieg DC, Cooper KF, Strich R. 2020. The extent of cyclin C promoter occupancy directs changes in stress-dependent transcription. J Biol Chem 295: 16280–16291.

Thoreen CC, Kang SA, Chang JW, Liu Q, Zhang J, Gao Y, Reichling LJ, Sim T, Sabatini DM, Gray NS. 2009. An ATP-competitive mammalian target of rapamycin inhibitor reveals rapamycin-resistant functions of mTORC1. J Biol Chem 284: 8023–8032.

Wang K, Yan R, Cooper KF, Strich R. 2015. Cyclin C mediates stress-induced mitochondrial fission and apoptosis. Mol Biol Cell 26: 1030–1043.

Wang XJ, Yu J, Wong SH, Cheng AS, Chan FK, Ng SS, Cho CH, Sung JJ, Wu WK. 2013. A novel crosstalk between two major protein degradation systems: regulation of proteasomal activity by autophagy. Autophagy 9: 1500–1508.

Westerling T, Kuuluvainen E, Makela TP. 2007. Cdk8 is essential for preimplantation mouse development. Mol Cell Biol 27: 6177–6182.

Yang S, Wang X, Contino G, Liesa M, Sahin E, Ying H, Bause A, Li Y, Stommel JM, Dell’antonio G et al. 2011. Pancreatic cancers require autophagy for tumor growth. Genes Dev 25: 717–729.

